# Future climate change-related decreases in food quality may affect juvenile Chinook salmon growth and survival

**DOI:** 10.1101/2022.08.28.505594

**Authors:** Jessica Garzke, Ian Forster, Caroline Graham, David Costalago, Brian P.V. Hunt

## Abstract

The global temperature increase due to global change is predicted to be between 3.3 – 5.7°C by 2100 leading to changes at the base of the marine food web in species composition, abundance, and quality at the base of the marine food web leading to flow-on effects of higher trophic levels such as fish and humans. Changes in marine prey availability and nutritional quality can affect juvenile salmon conditions (i.e., growth, condition, and mortality) during the early marine phase. There is limited knowledge of the interplay between prey availability and prey quality and the importance of food quality under food-satiated conditions. Here, a three-phase feeding experiment measured the effects of nutritional quality (fatty acid composition and ratios) on juvenile Chinook salmon (*Oncorhynchus tshawytscha*) condition. Experimental diets represented the present three different climate scenarios with a present-day diet (*Euphausia pacifica*), a control diet (commercial aquaculture diet), and a predicted IPCC worst-case scenario diet with low essential fatty acid concentrations (IPCC SSP5-8.5). We tested how potential future low quality food affects growth rates, body condition, fatty acid composition and mortality rates in juvenile Chinook salmon compared to present-quality prey. Fatty acids were incorporated into the salmon muscle at varying rates but, on average, reflected dietary concentrations. High dietary concentrations of DHA, EPA and high DHA:EPA ratios resulted in increased fish growth and condition. In contrast, low concentrations of DHA and EPA and low DHA:EPA ratios in the diets were not compensated for by increased food quantity. This result highlights the importance of considering food quality when assessing fish response to changing ocean conditions.

**Highlights:**

- Climate change may decrease the quality of salmon prey through changes in the fatty acid composition.
- Low dietary essential fatty acid levels reduce growth and condition and increase mortality rates in juvenile Chinook salmon.
- Food quality changes within zooplankton species but also by changes between species.
- Results suggest potential cascading effects on higher trophic levels when zooplankton species composition shifts to lower quality species.
- Higher food intake cannot compensate for low food quality.

## Introduction

Zooplankton play a fundamental role in aquatic food webs by consuming phytoplankton and bacteria and being the prey of higher trophic levels. At different life stages, many fish species derive their nutrition predominantly from zooplankton, which provide energy and essential biomolecules such as fatty and amino acids (Copeman and Laurel, 2010; Litz et al., 2017; Paulsen et al., 2014; Reitan et al., 1997). The nutritional quality of zooplankton prey directly affects fish condition, survival, and growth (Ahlgren et al., 2005; Daly et al., 2010; Olsen, 1999). Zooplankton nutritional quality varies significantly among species, and changes in the composition may therefore play an essential role in the feeding success, growth, and mortality of fish (Burke et al., 2013; Miller et al., 2014, 2013), particularly during vulnerable phases (Beamish et al., 2004; Pearcy and McKinnell, 2007). Understanding how prey composition impacts consumers has become a necessary step for estimating the survival of commercially important fish species.

Environmental variability in aquatic ecosystems is a key determinant of phenology, abundance, and species composition of zooplankton prey for foraging fish (Mackas et al., 2013; Mahara et al., 2019). This environmental variability occurs in space (e.g., bathymetry, tidal mixing) and time across a range of scales imposed by both natural cycles (e.g., seasonality, climate oscillations) and anthropogenic forcing (e.g., warming, acidification)(Vasseur and McCann, 2007). Over the past decades, the global mean surface temperature increased by ∼0.2°C per decade, reaching 1.0°C above the pre-industrial period in 2017 (Hoegh-Guldberg et al., 2019) and the newest predictions indicate a potential temperature increase of between 2.01 – 4.07°C of the global mean sea surface temperature (worst-case scenario; SSP5-8.5) by 2100 (IPCC, 2021). These temperature changes can lead to notable changes at the base of aquatic food webs through species-range shifts (Benedetti et al., 2021; McGinty et al., 2021; Poloczanska et al., 2013), shifts in phenology (Garzke et al., 2015; Parmesan and Yohe, 2003; Poloczanska et al., 2013; Walther et al., 2002), and reduced abundance and body size (Garzke et al., 2015).

Marine species have extended their range at an average rate of 72 km/decade towards the poles and into deeper waters responding to warmer environments (Lotze et al., 2010; Poloczanska et al., 2013). Specifically for marine zooplankton, time series analyses in the North Pacific and Atlantic Ocean showed that smaller bodied warmer-water species appeared and larger-bodied cold-water species disappeared (Beaugrand et al., 2002; Mackas et al., 2013, 2007; Preikshot et al., 2010)(Preikshot et al., 2010). Cold-water zooplankton species tend to have different life histories compared to their warmer-water species counterparts, as they have to diapause for overwintering and, therefore, store wax esters, triacylglycerols, phospholipids or diacylglycerol, such as euphausiids, *Calanus*, *Calanoides*, *Eucalanus*, *Neocalanus*, and *Rhinocalanus* whereas warm-water species have no energy-storage strategies and present a lower quality as a food item (Colombo et al., 2017; Hiltunen et al., 2021; Lee et al., 2006; Record et al., 2018). It is predicted that globally more non-native species will occur in new habitats. More native species will be extinct by 2050, and species turnovers of over 60% of the present biodiversity, implying direct effects on energetic pathways in marine food webs (Cheung et al., 2009; Heneghan et al., 2023). Indirect temperature-related changes, such as prey species composition, increased stratification, and corresponding changes in nutrient supply, may affect the production of essential biomolecules, such as fatty acids (FAs). These changes could alter the overall availability of fatty acids and the modification of an individual’s fatty acid profile (Garzke et al., 2016; Sahota et al., 2022; Stevens et al., 2022), ultimately affecting the efficiency of energy and biomass transfer through food webs (Pontavice et al., 2020; Pontavice, 2019; Ullah et al., 2018).

FA’s play a fundamental role in physiological processes in organisms from microorganisms to top predators, including humans, for energy storage, growth, and development, and trophic interactions in aquatic food webs (Dalsgaard et al., 2003; Parrish, 2009; Parrish et al., 2012). The precise composition of FAs is critical for the structure and function of all organisms and is thus considered an essential driver of organismal and ecosystem health and stability (Arts and Kohler, 2009; Parrish, 2009). Long-chain (i.e., >= 20 carbons) polyunsaturated fatty acids (LC-PUFA) and especially those of the n3 series, namely eicosapentaenoic acid (EPA; 20:5n-3) and docosahexaenoic acid (DHA; 22:6n-3), are key for brain and neural tissue growth, and cardiovascular health (Copeman et al., 2002). Despite their importance to animals, these n3 LC-PUFAs cannot be biosynthesized in sufficient quantity to meet their needs, and they must be obtained from their prey (Arts and Kohler, 2009; Hixson and Arts, 2016). LC-PUFAs are mainly synthesized by phytoplankton in aquatic food webs, especially by diatoms, cryptophytes, and dinoflagellates (Brett and Müller-Navarra, 1997). As primary consumers of phytoplankton, zooplankton act as conduits controlling the quantity and quality of LC-PUFA to higher trophic levels such as juvenile salmon. Accordingly, changes in phytoplankton FAs, zooplankton fatty acids, or the transfer between these two organisms will have cascading effects through the marine food web (Dalsgaard et al., 2003). Zooplankton consumers, such as juvenile fish, will be affected by the direct temperature-related impact on the organism’s metabolism, affecting their growth, reproduction, and survival, and the indirect temperature-related effects by changed bottom-up effects on the zooplanktons prey source, phytoplankton (Azani et al., 2021).

Global climate change could significantly affect the availability of long-chain omega-3 PUFAs in two not mutually exclusive ways: 1) by decreasing overall plankton biomass and growth rates and 2) by reducing the content of PUFAs, due to changes in zooplankton species composition with varying FA accumulation rates. For example, Hixson and Arts (2016) predicted that, in response to a 2.5°C increase in surface water temperature, phytoplankton could reduce DHA synthesis by up to 28% globally. Holm et al. (2022) predicted that EPA content in marine phyto- and zooplankton would substantially decline in the next century globally by 2%, ranging from 0.7% to 28.5%, depending on the geographic location. Since phytoplankton are the principal suppliers of DHA and EPA to higher trophic levels (e.g., fish), a reduction in the synthesis of DHA and EPA by phytoplankton as water temperatures rise has the potential to translate into a profound impact on the total supply of DHA and EPA to aquatic ecosystems, and hence the availability of these FAs to humans. Holm et al. (2022) even predicted that such a decline in essential FAs could lead to deleterious effects on fisheries.

In this study, we tested the climate-driven changes in zooplankton FA composition on fish condition, growth, mortality, and FA profiles. We performed a feeding experiment where juvenile Chinook salmon were fed either *Artemia* sp., which has an extremely low DHA and EPA concentration and low DHA:EPA ratio, a nutritionally enhanced control diet (commercial aquaculture), and *Euphausia pacifica* (a present-day diet representative).

## 2.1 Materials & Methods

### 2.1 Ethics Statement

All animals and procedures used in this experiment were approved by DFO Pacific Region - Animal Care Committee (AUP # 2017-008).

### 2.2 Experimental design

This study was conducted on the premises of Fisheries and Oceans Canada at the Pacific Science Enterprise Centre in West Vancouver, British Columbia, Canada. Juvenile Chinook salmon (ocean type, Harrison River strain) from the brood year 2018 (4.14 g ± 0.72 body weight; 73.6 mm ± 4.67 standard length) were obtained from Capilano River Hatchery (49.35° N 123.11° W) and transported to the research facilities. Before starting the trial, the fish were fed commercial salmon feed (Bio-Oregon, BioVita, Skretting Canada). The juvenile fish were acclimated in freshwater tanks for one week before transitioning to salt water, and only one individual died during the transition. Fish were not fed on the day of the transfer. The remaining 517 fish were used in the experiment (Fig. 1). The rearing system consisted of nine 200 L oval fibreglass tanks, each supplied with flow-through seawater (drawn from Burrard Inlet and passed through a sand filter) at 9 L min^-1^ and provided with supplemental air through an air stone. The overhead lights supplied lighting, and the photoperiod followed the natural day length. The fish were randomly assigned to the tanks, and each tank was assigned to three treatments in triplicate (n=3, Fig. 1). Fish were maintained in the experimental set-up for 36 days. The experimental phase was split into three individual phases (Fig. 1): (1) the acclimation phase when the fish were transferred to the experimental tanks (Phase 1); (2) weaning fish from control diets to their respective treatment diets (6 days; Phase 2); (3) feeding phase on experimental diets (29 days; Phase 3). Over the experimental period, the fish were held at an average temperature of 11.6⁰C (± 0.61⁰C).

**Fig. 1.**
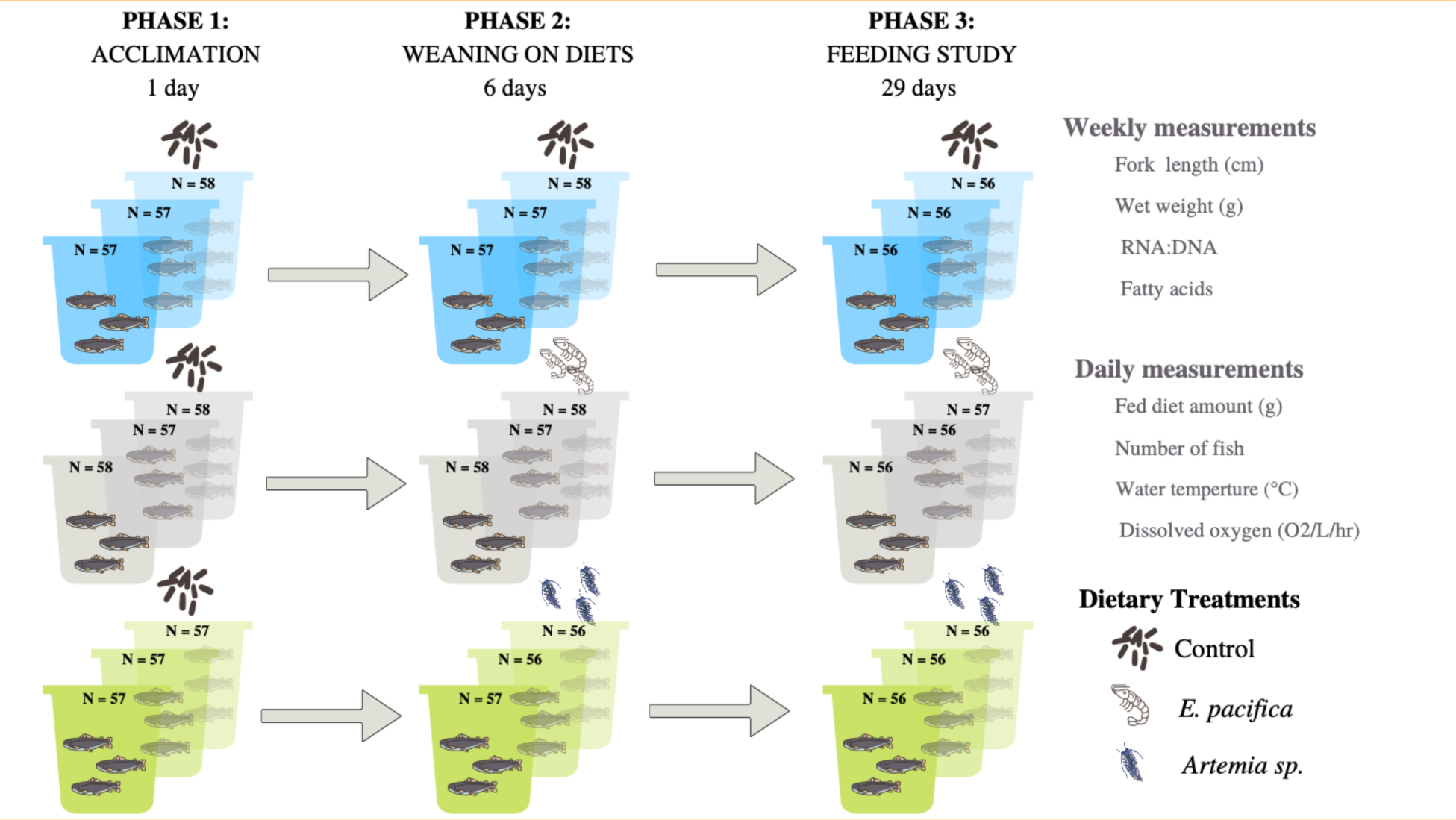
Three communities received a control diet (commercial aquaculture diet), three received *Euphausia pacifica* (present-day diet species), and three received *Artemia* sp. (climate change diet). Phase 1 lasted one day to acclimate fish to smaller tanks, and fish received the control diet; Phase 2 was six days to wean fish onto their respective diet treatments, and Phase 3 was the feeding study. Juveniles received their treatment diets until satiation.

### 2.3 Feeding

The three diet treatments were: 1) a commercial control (Bio-Oregon, BioVita, Skretting Canada), frozen *Euphausia pacifica* (Euphausia pacifica Canada) as present-day feeding condition representative, and frozen *Artemia* sp. as low-quality, climate change feeding condition representative (San Fransisco Bat Brand® Sally’s Frozen Spirulina Brine Shrimp™, Table 1). The commercial control had the lowest moisture content of the three diets (avg. = 5.79% ± 0.007), while *E. pacifica* and *Artemia* sp. had much higher moisture content (*E. pacifica*: avg. = 82.09% ± 0.24; *Artemia* sp.: 91.63% ± 0.32; Table 1, Fig. 1). The commercial control feed pellets, *Artemia* sp. and *E. pacifica* were thawed before feeding.

**Table 1.** Nutritional information of dietary treatments used in the experiment, for total fatty acids (TFA), saturated fatty acids (SFA), mono-unsaturated fatty acids (MUFA), polyunsaturated fatty acids (PUFA), docosahexaenoic acid (DHA), eicosatetraenoic acid (EPA), and DHA-to-EPA ratio (DHA:EPA). (DW: dry weight)

| Ingredients | Control | *Euphausia pacifica* | *Artemia* sp. |
| --- | --- | --- | --- |
| kJ g<sup>-1</sup> (DW) | 23.67 | 19.56 | 20.44 |
| TFA (g g<sup>-1</sup> DW) | 161.483 (± 11.112) | 24.069 (± 13.638) | 7.503 (± 4.118) |
| SFA (%) | 32.10 (± 0.10) | 28.47 (± 0.11) | 20.63 (± 0.10) |
| MUFA (%) | 29.32 (± 0.68) | 24.49 (± 0.22) | 30.34 (± 0.51) |
| PUFA (%) | 38.36 (± 1.85) | 46.81 (± 0.13) | 38.38 (± 0.42) |
| DHA (%) | 11.02 (± 0.58) | 16.80 (± 0.24) | 0.68 (± 0.94) |
| EPA (%) | 14.06 (± 0.28) | 16.11 (± 0.15) | 2.55 (± 0.15) |
| DHA:EPA | 0.78 (± 0.02) | 1.04 (± 0.02) | 0.28 (± 0.40) |

All diets were offered four times daily (7:30 – 8:00 am; 11:00 – 11:30 am; 12:30 – 1:00 pm; 2:00 – 2:30 pm) in smaller portions until the fish stopped actively feeding (*satiation*) depending on the individual needs of the treatment fish (Yu and Sinnhuber, 1979). The daily provided feed amounts were recorded. The tanks were cleaned three times a week after the last feeding by stopping the water flow and brushing the tank walls and bottom while the drain was plugged with a stopper. The sediment was then siphoned and cleaned. After the cleaning, the water flow was restarted, and the drain was unplugged.

### 2.4 Sampling and laboratory analyses

Throughout the experiment, four fish were randomly sampled from each tank weekly. Fish were removed from the tanks using nets and then euthanized in TMS^TM^ (0.4 g L^-1^). All fish were weighed to the nearest 0.1 g (Mettler Toledo EXCELLENCE Plus balance, model XP16001L), and fork length (FL) was measured (mm). Individual fish were placed into separate Whirlpak^TM^ bags, labelled with a unique identifier, and frozen at -80 °C. The time between the mortality of the fish and the fish being placed in the -80 °C freezer was less than two minutes. Tissue was sampled from each fish (n=336) for RNA:DNA ratios and fatty acids analysis using sterilized dissection tools. Before dissection, the dissection area was cleaned first with 95% ethanol and then with EZ-Zyme (Integra™ Miltex™ EZ-Zyme™ All-Purpose Enzyme Cleaner, Fisher Scientific, Cat No 12-460-427). White muscle tissue (400 – 600 mg) was collected close to the dorsal fin for RNA:DNA and fatty acid analysis. Tissue samples were then returned to a -80°C freezer until further analysis.

#### 2.4.1 RNA:DNA analysis

The ratio of RNA and DNA of juvenile Chinook was determined according to Garzke et al. (2022) using the cyanine base fluorescence dye RiboGreen^Ⓡ^ (Thermo Fisher Scientific). Nucleic acids were extracted from freeze-dried white muscle tissues, which had been homogenized in 1% N-lauroylsarcosine extraction buffer (VWR, Cat. No. VWRV0719-500G), the supernatant was diluted to match the RiboGreen saturation range (1:10, 1:20, 1:30, 1:50, 1:100). In short, we used a DNA standard (ranging from 0-3 µg mL^-1^; calf thymus, VWR, Supp. No. MB-102-01100) and RNA (ranging from 0-3 µg L^-1^; Escherichia *coli* 16S and 23S rRNA, Quant-iT RiboGreen RNA assay Kit Thermo Fisher Scientific Cat. No. R11490). Triplicate technical replicates from the diluted sample were added to a black, flat-bottom 96-well microtiter plate. RiboGreen^Ⓡ^ reagent was added to the homogenized muscle tissue samples and incubated in the dark for 5min; initial fluorescence (F_1_) readings were taken using a VarioSkan Flash Microplate Reader (ThermoFisher Scientific). RNase A (bovine pancreas, Alpha Aesar, Cat. No. CAAAJ62232-EX3) was added to each well and incubated at 37 °C for 30min in the dark, followed by a second fluorescence reading of the DNA only (F_2_). Finally, we calculated the RNA concentration using F_1_ - F_2_. RNA and DNA concentrations were calculated based on the standard curves, followed by calculations of RNA:DNA.

#### 2.4.2 Fulton K, growth rates, and mortality rates

For each sampling date and each diet treatment, we calculated the average body condition factor as Fulton’s K:

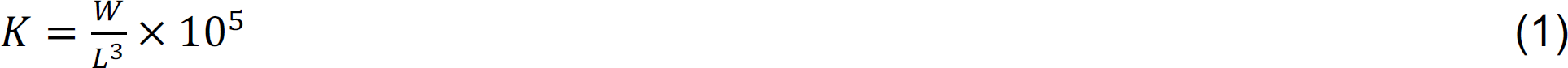

Where *W* is the wet body mass (g), and *L* is length (mm) (Fulton, 1904). Specific growth rates (SGR) for the weight (% W d^-1^) were measured for experimental

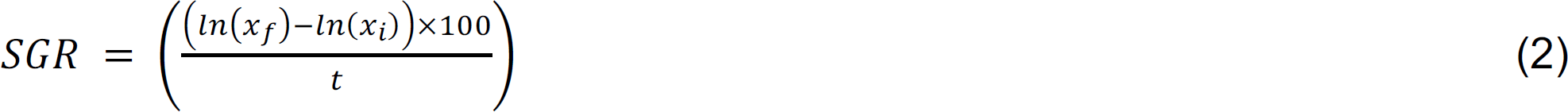

Where *X* is the weight of the last sampling day of Phase 3, *x_i_* is the weight of the first sampling day of Phase 3, and *t* is the time (days) between observations.

We calculated the instantaneous mortality rate (*Z*) by comparing the numbers of fish from one sampling day to the next for the duration of the experiment using (Sundby et al., 1989):

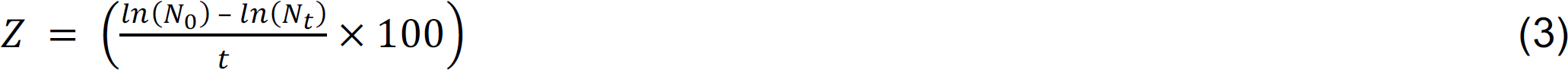

N_0_ is the initial number of fish, N_t_ is the number of remaining fish between sampling days, and *t* is the number of days between samplings.

#### 2.4.3 Fatty Acid analysis

FAs were analyzed using a modified protocol following Puttick et al. (2009) in a one-step fatty acid methyl ester (FAME) method. White muscle tissue samples were weighed (wet weight), freeze-dried, and the respective dry weights measured for calculations of moisture content. FAs were methylated with 3 N HCl in CH_3_OH (Sigma-Aldrich cat. #90964-500ML) with hexane (containing nonadecanoic acid C19:0 as an internal standard) at 80 C for 18 h. FAMEs were analyzed with a gas chromatograph (Scion 436-GC, Scion Instruments Canada, Edmonton, Alberta, Canada) using a 50 m column (CP Sil-88, Agilent Technologies, Santa Clara, California, USA) and a flame ionization detector (FID). Peaks were identified against external standards (GLC 455 and GLC 37 Nu-chek Prep, Inc., Elysian, Minnesota, USA) and measured before and after each sample measurement session.

All fatty acids were measured and reported as relative contents of total fatty acid methyl esters. Concentrations of 28 fatty acids were calculated using calibration curves of the known standard (C19:0). We calculated bioaccumulation factors (Kainz et al., 2006) of DHA, EPA, and ALA as

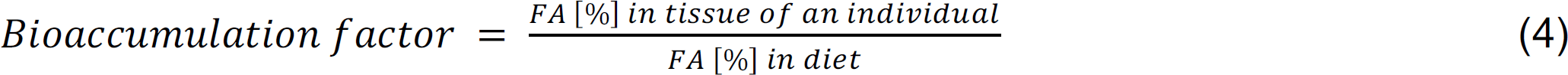

The bioaccumulation factor indicates consumers’ internal synthesis or retention of FAs. Specifically, the retention or internal synthesis of a FA increases with the increasing value of the bioaccumulation factor, and it is positive if the factor is >1.

### 2.5 Statistical analysis

We used linear mixed models repeated measures analysis to examine the effects of the three diet treatments and sampling date on fish condition (RNA:DNA, Fulton’s K) and individual fatty acid composition in fish muscle tissues across experimental Phase 3 to examine the effects of the three different prey qualities (using RStudio Version 1.1.456, lme4 R package (Bolker et al., 2009)). The tank number was set as a random factor.

Non-parametric PERMANOVA was used to test for differences in fish fatty acid composition between the three diet treatments. Subsequently, similarity percentages (SIMPER) were analyzed to identify the FA species that contributed most to the Bray-Curtis dissimilarities among the diet treatments. All multivariate tests were performed using the R package vegan (Oksanen et al., 2017).

Kolmogorov-Smirnov analysis was performed to test for normal distribution (α = 0.05). Analysis of Variance (ANOVA) was performed to identify differences between SGR between diet treatment, followed by a Tukey honest significance difference post hoc test. For non-normally distributed data, a Kruskal-Wallis test (Kassambara, 2020), followed by analyzing the effect size (Η^2^) and a pairwise comparison using the Dunn’s test were used to identify differences in instantaneous mortality rate (Z) and diet fatty acid data. The error rate for all tests was held at a 5% probability. All statistical tests were run at a significance threshold of α=0.05.

## 3. Results

Throughout the experiment, higher quantities of *Artemia* sp. were voluntarily eaten by fish compared to the control and *E. pacifica*. Percentage body weight (BW) consumed per day was: *Artemia* sp., 2.13% BW d^-1^ (of dry matter) ± 0.42 (range: 1.43% BW d^-1^ – 2.86% BW d^-1^); control, 1.49% BW d^-1^ (of dry weight) ± 0.34% BW (range: 0.99% BW d^-1^ – 2.35% BW d^-1^); *E. pacifica,* 1.68% BW d^-1^ (of dry matter) ± 0.66% (range: 1.04% BW d^-1^ – 2.50% BW d^-1^), Fig 2). Fish consuming *Artemia* sp. during Phase 3 had an average mortality rate of 1.96 ind d^-1^ (± 0.59) which was significantly higher than for fish consuming *E. pacifica* (1.21 ind d^-1^ ± 0.05; p = 0.04), and the control feed (1.23 ind d^-1^ ± 0.07; p = 0.04).

**Fig. 2.**
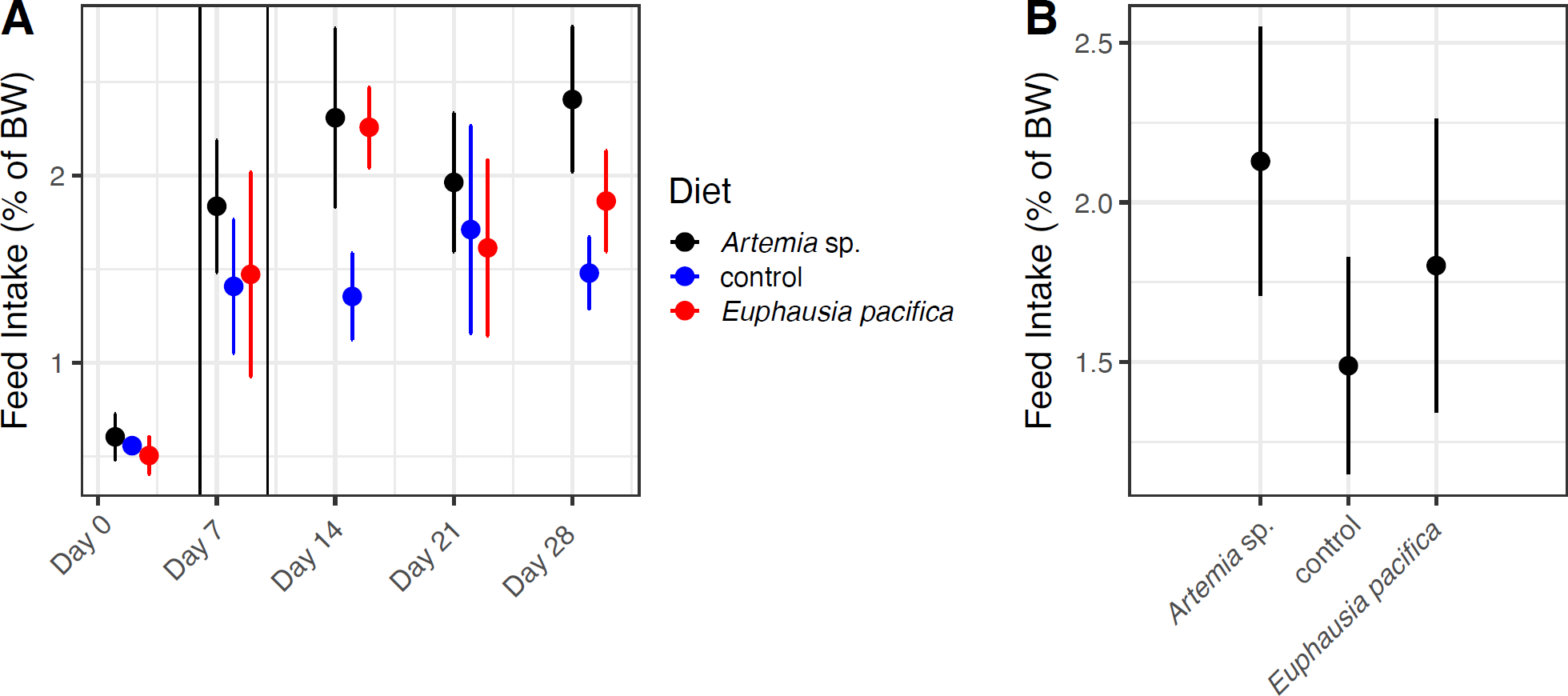
A) Voluntary feed intake (mean values (± Std Dev) by juvenile Chinook salmon as A) % of body weight dry matter over time, and B) average voluntary feed intake for each diet treatment during Phase 3.

### 3.1 Fish condition

Chinook salmon RNA:DNA ratios did not differ significantly among the three diet treatments in Phase 1 and Phase 2 (weaning) (Fig 3A, Table 2, Table S2). When fish transitioned from Phase 2 into Phase 3 (feeding study), within the first week, RNA:DNA of *E. pacifica* and control treatments increased by 7.47% and 21.21%, respectively, and *Artemia* sp. treatments RNA:DNA decreased by 47.38% (Fig 3A).

**Fig. 3.**
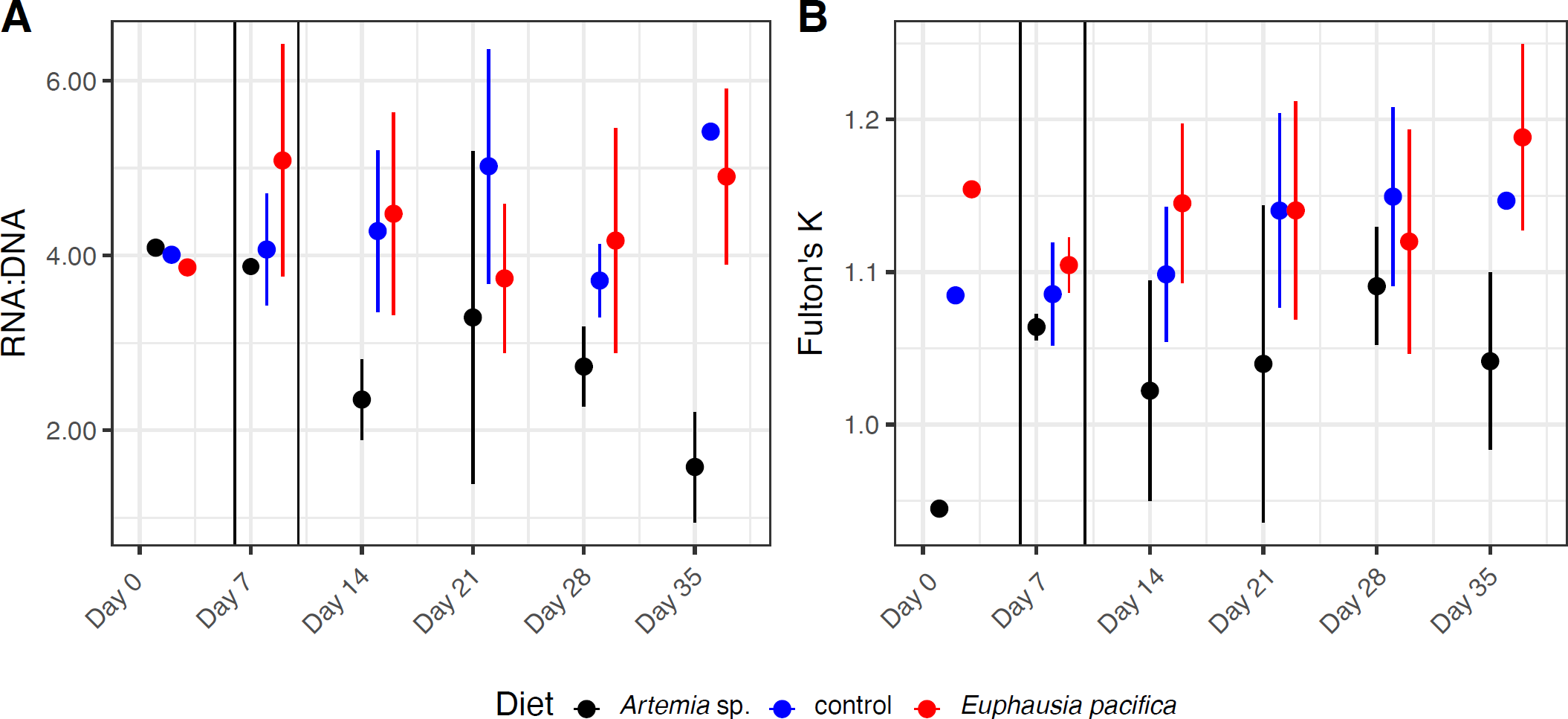
Mean values (± Std Dev) of A) RNA:DNA ratios and B) Fulton’s K condition index across the experimental period.

**Table 2.** Analysis of deviance of linear mixed models based on full models testing effects of diet treatments over time on juvenile Chinook with the tank as random effects (full linear mixed model results see Table S2. TFA = Total fatty acids, SFA = saturated fatty acids, MUFA = monounsaturated fatty acids, PUFA = polyunsaturated fatty acids, EFA = essential fatty acids, DHA = docosahexaenoic acid, EPA = eicosapentaenoic acid, ALA = alpha-linolenic acid.

|  | Predictor | Chi<sup>2</sup> | Df | P value |
| --- | --- | --- | --- | --- |
| RNA:DNA | Treatment | 77.194 | 2 | <0.001 |
|  | Sampling date | 10.989 | 1 | <0.001 |
| Fulton K | Treatment | 16.959 | 2 | <0.001 |
|  | Sampling date | 24.902 | 1 | <0.001 |
| TFA | Treatment | 5.168 | 2 | 0.075 |
|  | Sampling date | 14.318 | 1 | <0.001 |
| SFA | Treatment | 37.997 | 2 | <0.001 |
|  | Sampling date | 84.812 | 1 | <0.001 |
| MUFA | Treatment | 15.099 | 2 | <0.001 |
|  | Sampling date | 0.642 | 1 | 0.422 |
| PUFA | Treatment | 28.114 | 2 | <0.001 |
|  | Sampling date | 15.279 | 1 | <0.001 |
| EFA | Treatment | 30.242 | 2 | <0.001 |
|  | Sampling date | 18.258 | 1 | <0.001 |
| DHA | Treatment | 53.184 | 2 | <0.001 |
|  | Sampling date | 23.060 | 1 | <0.001 |
| EPA | Treatment | 29.836 | 2 | <0.001 |
|  | Sampling date | 58.886 | 1 | <0.001 |
| DHA:EPA | Treatment | 37.184 | 2 | <0.001 |
|  | Sampling date | 40.481 | 1 | <0.001 |
| ALA | Treatment | 244.25 | 2 | <0.001 |
|  | Sampling date | 2.84 | 1 | 0.0919 |

Throughout Phase 3, *Artemia* sp. treatments had a significantly lower RNA:DNA ratio (2.68 ± 1.1) than *E. pacifica* treatments (4.43 ± 0.94) and control treatments (4.78 ± 1.11). The RNA:DNA of the *Artemia* sp. treatment declined throughout Phase 3. The *E. pacifica* and control treatments had the highest RNA:DNA ratios and were not significantly different from each other (Table 2, Table S2).

Fulton’s K body condition index increased significantly after Phase 2 (Fig. 3B, Table 2). However, this increase was dependent on diet treatment. Fulton’s K during Phases 1 and 2 did not differ within or among diet treatments (Fig 3B). When the fish transitioned from Phase 1 into Phase 2, only minor changes in Fulton K were measured (*Artemia sp.*: -1.86%; *E. pacifica*: +1.88%; control: -3.6%). After the transition to Phase 3, after the first week, Fulton’s K increased by 5.61% and 2.78% in the *E. pacifica* and control treatments, respectively. Throughout Phase 3, Fulton’s K increased significantly in the *E. pacifica* and control treatments (start: control 1.11 ± 0.04, *E. pacifica* 1.12 ± 0.04; end: control 1.15 ± 0.05, *E. pacifica* 1.14 ± 0.05). Fulton’s K of *Artemia* sp. treatments was significantly lower than *E. pacifica* and control treatments.

Specific growth rates across Phase 3 were not significantly different among the three diet treatments (Table 3). The average weight increase of fish from the end of Phase 2 and the end of Phase 3 was 1.29% d^-1^ ± 0.40 for *Artemia sp.*, 1.35% d^-1^ ± 0.29 for control, and 1.91% d^-1^ ± 0.58 for *E. pacifica* (Fig 4A). Instantaneous mortality rate (Z) did not differ between treatments within experimental feeding Phase 3 (p = 0.57) but was highest for *Artemia* sp. treatments (∼1.7 vs. ∼1.2 for *E. pacifica* and control treatments; Fig 4B). The cumulative mortality was higher in Artemia treatments compared to *E. pacifica* and control treatments (Fig. S1)

**Table 3.** Analysis of Variance (ANOVA) results of specific growth rates (SGR) of fish from last day Phase 2 to last day of Phase 3; Df = degrees of freedom, Sum Sq = Sum of Squares, Mean Sq = Mean squares).

|  |  | df | Sum Sq | Mean Sq | F value | p |
| --- | --- | --- | --- | --- | --- | --- |
| SGR weight | Treatment | 2 | 0.8914 | 0.4457 | 3.202 | 0.113 |
|  | Residuals | 6 | 0.351 | 0.1392 |  |  |

**Fig. 4.**
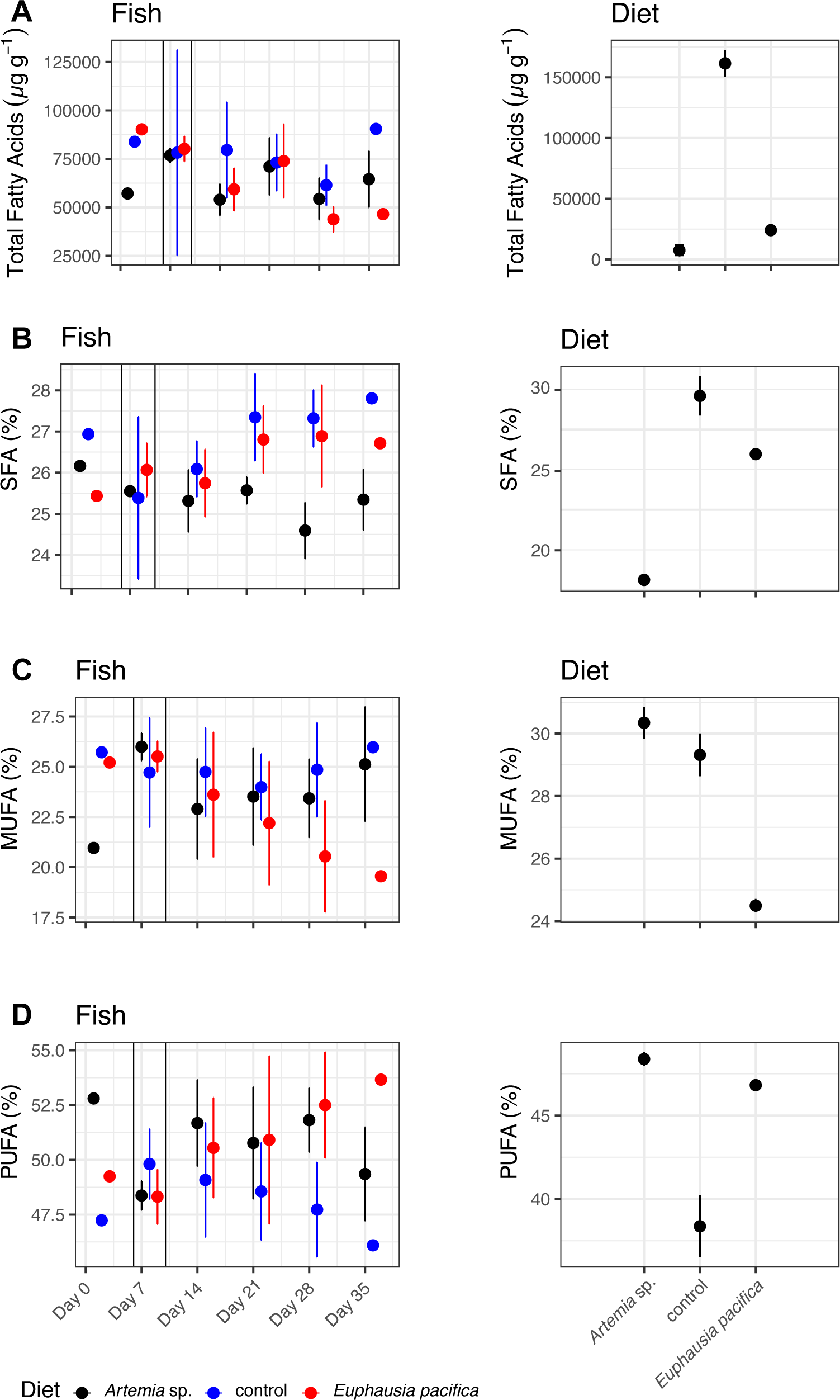
Fatty acid composition (mean values ± Std Dev) of Chinook white muscle tissue and diets in each treatment. A) Total fatty acids (TFA), B) saturated fatty acids (SFA), C) monounsaturated fatty acids (MUFA), and D) polyunsaturated fatty acids (PUFA). Mean values.

### 3.2 Fatty Acids (FA)

The PERMANOVA showed that fish FA profiles were significantly affected by diet during experimental Phase 3 (p <0.001; Table S3). DHA contributed the most (>20%) to the dissimilarity between all treatment pairs and was most remarkable for *E. pacifica* vs control (32%; Table S4). The proportion of ALA in *Artemia sp.* differed by 18% and 18.8% from *E. pacifica* and control, respectively, while oleic acid (C18:1n-9) differed by >12% between *E. pacifica* and control (Table S4). C16:0 was also detected as a contributor to the dissimilarity between control and *E. pacifica* and *Artemia* sp. We excluded this from our further analysis as C16:0 is a typical trophic marker for a fish meal, the main ingredient in the control feed. The total fatty acid concentration in fish did not differ significantly between the three diet treatments (Fig 4A, Table 2, Table S2).

#### 3.2.1 Fatty acid groups

Fish SFA percentages differed among the three diet treatments (Table 2, Table S2). SFA percentages were significantly higher in *E. pacifica* and control treatments compared to *Artemia* sp. treatments (Fig 4B, Table 2, Table S2). SFA percentage increased significantly from the first day of Phase 3 to the end of Phase 3 in *E. pacifica* and control treatments (*E. pacifica*: start 26.75% ± 0.92, end 26.82% ± 0.78; control: start 26.39% ± 0.95, end 27.46% ± 0.77 Fig 4B; Table 2, Table S2). The SFA percentage of fish in *Artemia* sp. treatments decreased in the same phase from 26.02% (± 0.81) to 25.47% (± 0.36) (Fig 4C). No significant difference was detected in the MUFA percentage between the different diet treatments (Fig 4C, Table 2, Table S2). PUFA contribution was significantly lower in control treatments (49.79% ± 2.11) compared to *Artemia* sp. (51.37% ± 2.52) and *E. pacifica treatments* (50.67% ± 3.24; Table 3). PUFA composition remained constant in *Artemia* sp., and *E. pacifica* treatments provided fish during Phase 3 (*E. pacifica*: start - 50.19% ± 3.03, end - 50.16% ± 3.08; *Artemia*: start - 50.92 ± 1.85, end - 51.50% ± 3.00; Fig 4D). PUFAs in the control treatment fish decreased slightly from the beginning to the end of Phase 3 (49.61% ± 2.01 - 48.60% ± 2.62).

DHA was significantly higher in *E. pacifica* treatments compared to control and *Artemia* sp. treatments (Table 2, Table S2), whereas DHA in control and *Artemia* sp. treatments were similar (Fig 5A). All fish had a comparable inter-treatment DHA percentage in Phase 2 (*Artemia sp.*: 30.68% ± 2.59; control: 29.25% ± 3.24; *E. pacifica*: 30.27% ± 1.87) before the fish were switched to their experimental diets. One week after the diet switch, all treatments slightly increased their DHA (*Artemia sp.*: +0.74%; control: 0.94%; *E. pacifica*: +1.59%). Over Phase 3, DHA slightly increased in the *E. pacifica* treatment (start 31.86% ± 4.07, end 32.42% ± 4.14), whereas DHA in control and *Artemia sp.* treatments slightly decreased over time (control: start 30.19% ± 2.24, end 29.37% ± 3.91; *Artemia sp.*: start 31.42% ± 2.97, end 27.95%± 4.4; Fig 5A).

**Fig. 5.**
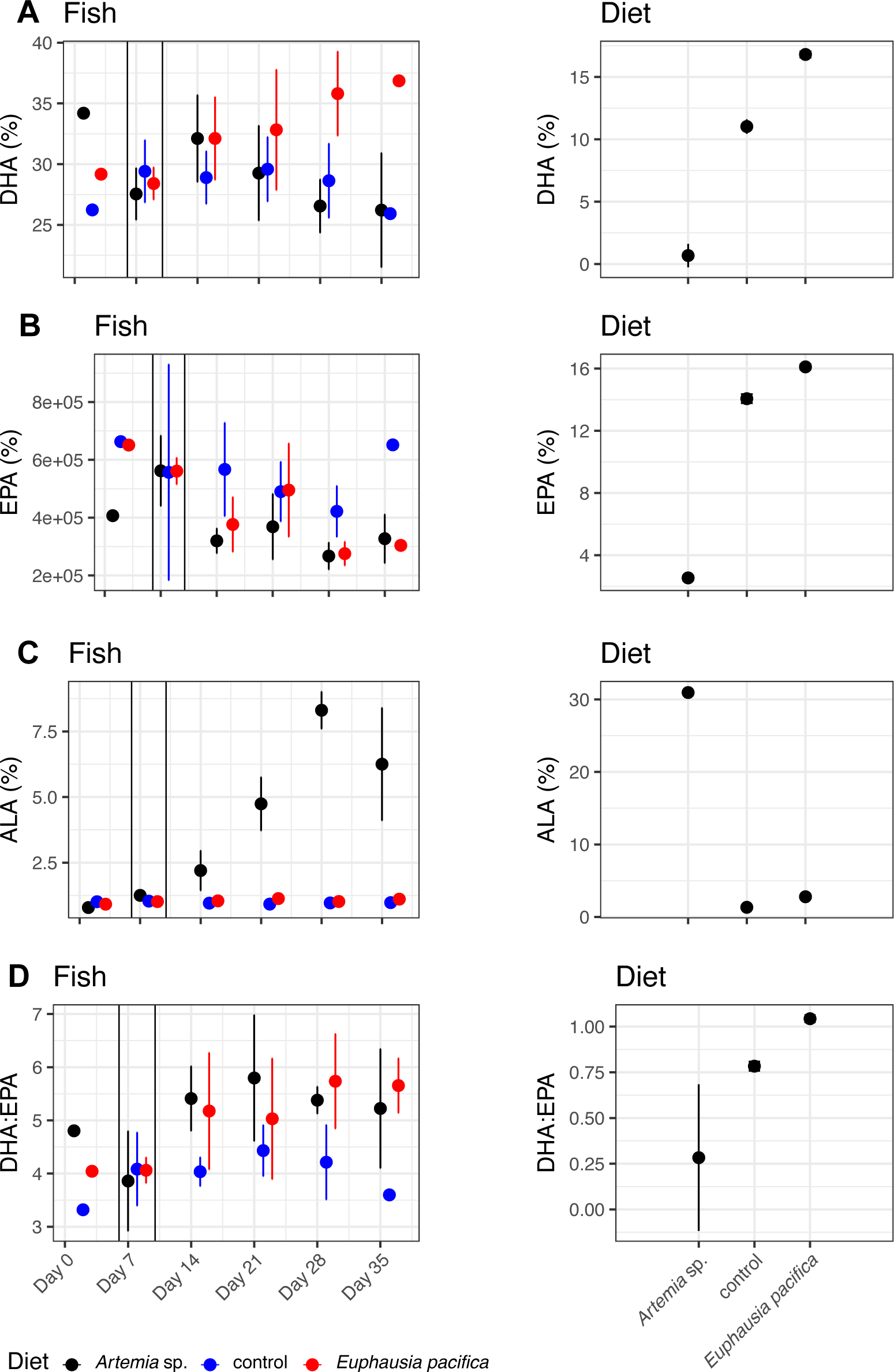
Mean fatty acid percentual composition (± Std Dev) compared to TFA in Chinook white muscle tissue and provided diet treatments. A) DHA, B) EPA, C), ALA, D) DHA:EPA.

During Phase 3, EPA decreased in *Artemia* sp. treatments (0.81%), control treatments (0.18%), and *E. pacifica* treatments (0.47%) within one week (Fig. 5B). The EPA percentage in *Artemia* sp. treatments decreased slightly over Phase 3, from 5.89% ± 0.31 to 4.93% ± 0.30, whereas in the *E. pacifica* and control treatments, EPA was constant (*E. pacifica*: start 6.40% ± 0.51, end 6.64% ±0.63; control: start 6.94% ±0.60, end 7.10% ±0.61; Fig. 5B).

ALA percentage was significantly higher in the *Artemia* sp. treatment compared to *E. pacifica* and control treatments (Table 2, Table S2; Fig. 5C). During Phase 2, ALA in all treatments was similar (*Artemia sp.*: 1.27% ± 0.07; control: 0.99% ± 0.15; *E. pacifica*: 0.99% ± 0.07). ALA increased in *Artemia* sp. after the complete diet switch at the beginning of Phase 3 (2.27%), but *E. pacifica* and control treatments remained relatively constant. ALA percentage increased over time during Phase 3 in *Artemia* sp. treatments (start 2.27% ± 0.65, end 6.72% ± 1.46). In *E. pacifica* treatments, the ALA percentage increased slightly over time (start 1.03% ± 0.10, end 1.25% ± 0.15), and in control treatments, the ALA percentage remained constant over time (start 0.91% ± 0.81, end 0.91% ± 0.11; Fig 5C).

DHA:EPA ratios were significantly affected by time (Table 2, Table S2). During Phase 2, DHA:EPA ratios were comparable among treatments (*Artemia sp.*: 4.64 ± 0.74; control: 4.18 ± 0.83; *E. pacifica*: 4.42 ± 0.41; Fig. 5D). After transitioning into Phase 3, DHA:EPA ratios increased in *Artemia* sp. and *E. pacifica* treatments (+0.61) (Fig 5D). The DHA:EPA ratio in the *Artemia* sp. and *E. pacifica* treatments increased slightly over experimental Phase 3 (Fig 5D). DHA:EPA ratios in the control treatment were significantly lower than *Artemia* sp. and *E. pacifica* and decreased slightly over Phase 3 (4.37 ± 0.47 - 4.19 ± 0.85). DHA:EPA ratios were highest in *Artemia*-fed Chinook in Phase 3 and slightly increased over time (start 5.34 ± 0.61, end 5.72 ± 1.21). *E. pacifica* treatments had a slightly lower DHA:EPA ratio than the *Artemia* sp. treatment (start 5.04 ± 0.98, end 4.96 ± 1.01).

We found that fish did not bioaccumulate ALA (all prey treatments) and EPA (control and *E. pacifica*), as the percentage of these fatty acids was lower in the tissues of the fish than in their diets (Table 4). In contrast, fish in all treatments bioaccumulated DHA, and the bioaccumulation factor was significantly higher in DHA deficient *Artemia* sp. diet than in the control or *E. pacifica* treatments. Despite the higher bioaccumulation, the relative concentration of DHA was significantly lower in muscle tissue of DHA-deprived *Artemia* sp. treatments than in the control and *E. pacifica* treatments.

**Table 4.** Mean bioaccumulation (± SD) of DHA, EPA and ALA across all three diet treatments over the experimental phase Phase 3.

|  | *Artemia* sp. | Control | *Euphausia pacifica* |
| --- | --- | --- | --- |
| **DHA** | **40.894** (± 7.460) | **2.874** (± 0.356) | **1.945** (±0.255) |
| **EPA** | **1.956** (± 0.122) | **0.486** (± 0.031) | **0.412** (± 0.039) |
| **ALA** | **0.212** (±0.053) | **0.296** (± 0.039) | **0.432** (± 0.062) |

## 4. Discussion

Our results show that low dietary concentrations of DHA and EPA and low DHA:EPA ratios induced by experimental treatment resulted in reduced growth and condition compared to a control diet and the present dominant prey species, *Euphausia pacifica* (Duguid, 2021). Furthermore, low concentrations of DHA and EPA and low DHA:EPA ratios were not compensated for by higher food quantity intake. This indicates that food quality needs to be considered when assessing fish response to changing ocean conditions. Below we evaluate the response of juvenile Chinook salmon to individual fatty composition as measured by condition factor, RNA:DNA ratio and growth rate. Finally, we assess the implications for the response of fish to climate-driven changes in zooplankton communities.

Dietary DHA:EPA ratios >2 are considered optimal for larval and juvenile marine fish (Bell et al., 1995; Watanabe, 1993), and ratios of ∼1 are sufficient in many species (Copeman and Laurel, 2010; Rodriguez et al., 1997). *E. pacifica* was ∼1, the control diet ∼0.75, and *Artemia* sp. below with ∼0.25 (representing diets such as crab larvae and polychaetes; Hiltunen et al. (2021)). Based on data for chum, coho, and Atlantic salmon, SFA and MUFA in diets should be approximately 33% (Turchini et al., 2009). All diets in the present study were below 33% in SFA and MUFA, respectively. Additionally, essential fatty acids, the sum of DHA, EPA, ALA, and linolenic acid, should contribute >1-2% of the diets’ dry weight (Tocher, 2010). In the present study, the contribution of EFA was highest in *Artemia* sp. and lowest in *E. pacifica* and the commercial control. Still, the individual FA composition varied significantly among feeds. *Artemia* sp. had the highest ALA concentration, contributing to the significantly higher amount of EFAs than *E. pacifica* and control diets, which had lower ALA concentrations. SFA was significantly lower in *Artemia* sp. than in both *E. pacifica* and control diets. These differences were related to specific growth and FA composition responses in the experimental fish.

The bioaccumulation factor of DHA in all diet treatments was higher than one and about 40 times higher in salmon fed with DHA-deprived *Artemia* sp. than the control and the *E. pacifica* fish. In contrast, differences in EPA and ALA bioaccumulation factors between the diet treatments were not retained in the salmon tissues; only the bioaccumulation factor for EPA fed with *Artemia* sp. was ∼1.9. This suggests that dietary EPA and ALA were not retained in the salmon tissues but instead likely used for cell energy synthesis by fish (Kainz et al., 2006; Závorka et al., 2021). Despite the high bioaccumulation factor, DHA was low in salmon of *Artemia* treatments, associated with lower growth rates, low body condition (RNA:DNA ratios, Fulton’s K), and high mortality rates. This finding corresponds with previous studies and demonstrates that maintaining a similar DHA content as the control group is costly (Závorka et al., 2021; Taipale et al., 2018; Murray et al., 2014). Závorka et al. (2021) showed that juvenile salmon fed under n-3 PUFA-deprived conditions had lower mitochondrial efficiencies of ATP production in muscle tissues. The fastest metabolic pathway synthesizing adenosine triphosphate (ATP) is from SFA, followed by MUFA, and PUFA by β-oxidation in the peroxisome (Turchini and Francis, 2009). In salmon, EPA is extensively used for β-oxidation when supplied in dietary surplus (Codabaccus et al., 2011; Stubhaug et al., 2005), and DHA is more conserved, irrespective of its dietary concentration (Tocher, 2010). Chinook SFAs in the *Artemia* treatments decreased over time, suggesting that this was related to the low nutritional SFA requiring ATP synthesis from stored SFA. Conversely, SFAs in *E. pacifica* and control treatments increased over the same experimental phase, indicating a sufficient supply of ATP for growth. We observed that Artemia-fed fish had higher intake rates than the control and *E. pacifica*-fed fish, which might have been a compensation mechanism leading to a higher energy intake to cover higher energy expenses.

The effects of food quality on fish growth demonstrated here and in other studies (Brett and Müller-Navarra, 1997; Malzahn et al., 2007b) highlight the important role of prey quality in fish condition and how changes in zooplankton due to climate change could affect zooplanktivorous consumers. In the case of zooplanktivorous fish, the quality of zooplankton prey can be determined by 1) the species composition of the zooplankton themselves or 2) by shifts within the zooplankton species due to changes in their phytoplankton prey. The fatty acid composition of zooplankton in the Salish Sea varies substantially among species (Costalago et al., 2020; Hiltunen et al., 2021). The relative proportion of these prey taxa can vary with ocean conditions, including whole community shifts from lipid-rich boreal species during cold conditions to lipid-poor species, including more gelatinous plankton, during warm conditions (El-Sabaawi et al., 2009a; Mackas et al., 2013). Such environmentally-driven shifts have been demonstrated to occur over decadal scales and in rapid response to recent marine heatwaves (Fisher et al., 2015; Miller et al., 2017; Peterson et al., 2017; Young et al., 2019). While distributional shifts in zooplankton are expected to be important, changes in their phytoplankton food base have also been demonstrated to be an important factor in their FA composition (El-Sabaawi et al., 2009a, 2009b). EFAs are primarily synthesized by phytoplankton and transferred through the food web via zooplankton to higher trophic levels. Phytoplankton have taxon-specific FA profiles; for example, diatoms have higher proportions of EPA than DHA, while dinoflagellates have higher proportions of DHA than EPA (Dalsgaard et al., 2003).

Fatty acid composition within species can also be affected by environmental conditions. For example, experimental studies showed that nitrogen and phosphorus limitation could increase TFA, SFA and n-3 and n-6 FA concentrations in phytoplankton (Bi et al., 2018; Malzahn et al., 2007a; Schoo et al., 2014); warming can increase the SFA to PUFA ratio (Hixson and Arts, 2016; Jin et al., 2020); and ocean acidification leads to reduced concentrations of TFA and PUFA (Meyers et al., 2019), but also that changes are species-specific and change with community species composition (Dörner et al., 2020; Leu et al., 2013). Such changes in nutritional quality at the base of the food web have been demonstrated to transfer to higher trophic levels, affecting consumers, e.g., when phytoplankton has higher concentrations of DHA and EPA, the same can be measured in zooplankton and secondary consumers show increased nutritional condition (RNA/DNA) (Malzahn et al., 2007a).

## 5. Conclusions

This study found that high dietary concentrations of DHA and EPA and high DHA:EPA ratios promoted the growth and condition of juvenile Chinook salmon. Furthermore, poor food quality, characterized by low concentrations of DHA and EPA and low DHA:EPA ratios, was not compensated for by increased food quantity. Such changes in food quality are potentially an important mechanism behind the response of zooplanktivorous fish to climate change. This highlights the importance of considering food quality when assessing fish response to changing ocean conditions. We recommend that future studies consider food quality when determining foraging conditions for juvenile salmon. Furthermore, species-specific data on zooplankton nutrition could be used to qualify zooplankton time series retrospectively to assess how foraging conditions have changed over time. Additionally, testing only various DHA:EPA ratios or concentrations with comparable base feed would be beneficial to understand the effect on juvenile Chinook growth and condition.

## Acknowledgements

We gratefully acknowledge the contributions of Lauren Porter and Katarina Doughty. L. Porter provided technical assistance and supported the experiment during sampling, feeding, and dissections. K. Doughty assisted with the fatty acid analyses, the experimental setup, and fish acclimation phase. Their work was essential to the quality of the results presented in this paper.

## Author contributions

Conceptualization: J.G, I.F, D.C., B.P.V.H.; Methodology: J.G., C.G.; Validation: J.G.; Data curation: J.G, C.G.; Writing – original draft: J.G.; Writing-review & editing J.G, I.F., C.G, D.C., B.P.V.H.; Supervision: B.P.V.H., I.F.; Project administration: B.P.V.H., I.F.; Funding acquisition: B.V.P.H., I.F.

## Funding

This study was supported by a Fisheries and Oceans Canada Strategic Program for Ecosystem-Based Research and Advice (SPERA) grant. The Tula-Mitacs Canada Grants IT09911 and IT13677 supported J. Garzke.

## Supplementary material

**Table S1.**
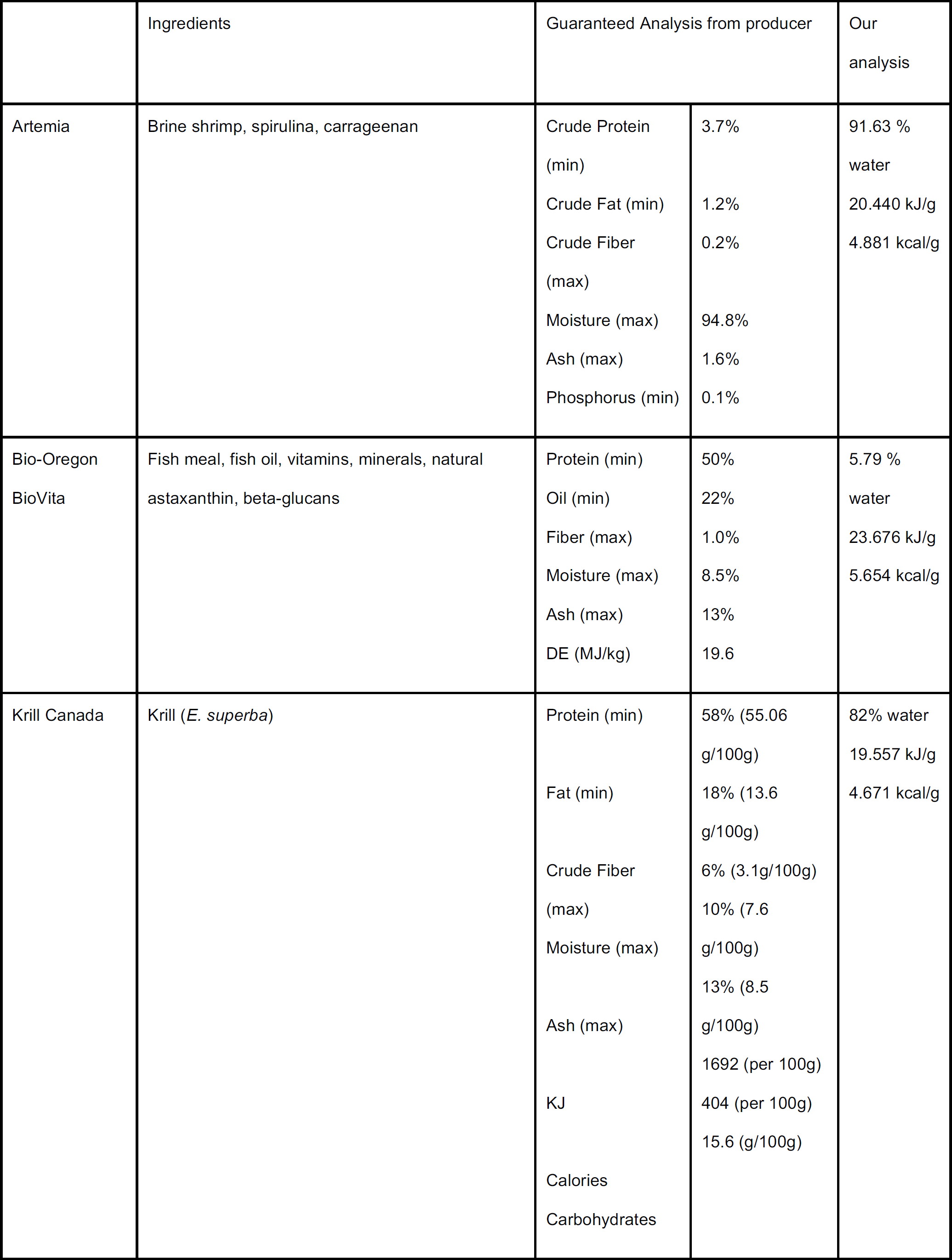
Feed composition (producer)

|  | Ingredients | Guaranteed Analysis from producer |  | Our analysis |
| --- | --- | --- | --- | --- |
| Artemia | Brine shrimp, spirulina, carrageenan | Crude Protein (min) | 3.7% | 91.63 % water |
|  |  | Crude Fat (min) | 1.2% | 20.440 kJ/g |
|  |  | Crude Fiber (max) | 0.2% | 4.881 kcal/g |
|  |  | Moisture (max) | 94.8% |  |
|  |  | Ash (max) | 1.6% |  |
|  |  | Phosphorus (min) | 0.1% |  |
| Bio-Oregon<br>BioVita | Fish meal, fish oil, vitamins, minerals, natural astaxanthin, beta-glucans | Protein (min) | 50% | 5.79 % water |
|  |  | Oil (min) | 22% | 23.676 kJ/g |
|  |  | Fiber (max) | 1.0% | 5.654 kcal/g |
|  |  | Moisture (max) | 8.5% |  |
|  |  | Ash (max) | 13% |  |
|  |  | DE (MJ/kg) | 19.6 |  |
| Krill Canada | Krill (*E. superba*) | Protein (min) | 58% (55.06 g/100g) | 82% water |
|  |  | Fat (min) | 18% (13.6 g/100g) | 19.557 kJ/g |
|  |  | Crude Fiber (max) | 6% (3.1g/100g) | 4.671 kcal/g |
|  |  | Moisture (max) | 10% (7.6 g/100g) |  |
|  |  | Ash (max) | 13% (8.5 g/100g) |  |
|  |  | KJ | 1692 (per 100g) |  |
|  |  | Calories | 404 (per 100g) |  |
|  |  | Carbohydrates | 15.6 (g/100g) |  |

**Table S2.** Full statistical analysis of linear mixed models. TFA = Total fatty acids, SFA = saturated fatty acids, MUFA = monounsaturated fatty acids, PUFA = polyunsaturated fatty acids, EFA = essential fatty acids, DHA = docosahexaenoic acid, EPA = eicosapentaenoic acid, ALA = alpha-linolenic acid

|  |  | Predictor |  |  |  | Random Effects |  |  | N | Marginal R<sup>2</sup> |
| --- | --- | --- | --- | --- | --- | --- | --- | --- | --- | --- |
| | | Intercept | Control | Krill | Sampling date | $\sigma^2$ | t<sub>Tank</sub> | N<sub>Tank</sub> | | |
| RNA/DNA | Estimates | 261.77 | 1.27 | 1.21 | -0.01 | 1.19 | 0.00 | 9 | 254 | 0.268 |
|  | CI | 145.08 – 378.46 | 0.86 – 1.68 | 0.79 – 1.62 | -0.02 – -0.01 |  |  |  |  |  |
|  | p-value | **<0.001** | **<0.001** | **<0.001** | **<0.001** |  |  |  |  |  |
|  | df | 249.21 | 5.99 | 5.72 | 249.22 |  |  |  |  |  |
| Fulton K | Estimates | -19.06 | 0.05 | 0.03 | 0.00 | 0.00 | 0.00 | 9 | 204 | 0.168 |
|  | CI | -27.08 – -11.03 | 0.02 – 0.08 | 0.00 – 0.06 | 0.00 – 0.00 |  |  |  |  |  |
|  | p-value | **<0.001** | **0.007** | **0.041** | **<0.001** |  |  |  |  |  |
|  | df | 200.00 | 5.98 | 5.09 | 200.00 |  |  |  |  |  |
| TFA | Estimates | 5483.10 | 5.63 | -4.07 | -0.31 | 731.01 | 2.26 | 9 | 2.64 | 0.072 |
|  | CI | 2652.09 – 8314.10 | -4.86 – 16.11 | -14.57 – 6.44 | -0.47 – -0.15 |  |  |  |  |  |
|  | p-value | **<0.001** | 0.238 | 0.376 | **<0.001** |  |  |  |  |  |
|  | df | 258.58 | 6.10 | 5.64 | 258.59 |  |  |  |  |  |
| SFA | Estimates | -414.27 | 0.97 | 0.65 | 0.02 | 0.81 | 0.01 | 9 | 264 | 0.340 |
|  | CI | -508.71 – -319.82 | 0.58 – 1.36 | 0.26 – 1.05 | 0.02 – 0.03 |  |  |  |  |  |
|  | p-value | **<0.001** | **0.001** | **0.007** | **<0.001** |  |  |  |  |  |
|  | df | 257.96 | 6.09 | 5.74 | 257.97 |  |  |  |  |  |
| MUFA | Estimates | -86.86 | 0.96 | -0.55 | 0.01 | 6.76 | 0.00 | 9 | 264 | 0.056 |
|  | CI | -359.05 – 185.27 | -0.00 – 1.93 | -1.52 – 0.41 | -0.01 – 0.02 |  |  |  |  |  |
|  | p-value | 0.530 | 0.051 | 0.208 | 0.426 |  |  |  |  |  |
|  | df | 258.80 | 6.11 | 5.60 | 258.81 |  |  |  |  |  |
| PUFA | Estimates | 605.93 | -1.92 | -0.08 | -0.03 | 7.21 | 0.00 | 9 | 264 | 0.139 |
|  | CI | 324.95 – 886.92 | -2.92 – -0.92 | -1.08 – 0.92 | -0.05 – -0.02 |  |  |  |  |  |
|  | p-value | **<0.001** | **0.003** | 0.856 | **<0.001** |  |  |  |  |  |
|  | df | 258.80 | 6.11 | 5.60 | 258.81 |  |  |  |  |  |
| EFA | Estimates | 685.73 | -1.82 | 0.41 | -0.04 | 803 | 0.00 | 9 | 264 | 0.152 |
|  | CI | 389.06 – 982.40 | -2.87 – -0.77 | -0.64 – 1.46 | -0.05 – -0.02 |  |  |  |  |  |
|  | p-value | **<0.001** | **0.005** | 0.375 | **<0.001** |  |  |  |  |  |
|  | df | 258.80 | 6.11 | 5.60 | 258.81 |  |  |  |  |  |
| DHA | Estimates | 1004.45 | 0.09 | 3.67 | -0.06 | 14.73 | 0.00 | 9 | 264 | 0.220 |
|  | CI | 602.68 – 1406.21 | -1.33 – 1.52 | 2.24 – 5.10 | -0.08 – -0.03 |  |  |  |  |  |
|  | p-value | **<0.001** | 0.880 | **0.001** | **<0.001** |  |  |  |  |  |
|  | df | 258.80 | 6.11 | 5.60 | 258.81 |  |  |  |  |  |
| EPA | Estimates | -375.81 | 0.71 | 0.05 | 0.02 | 0.89 | 0.00 | 9 | 2.64 | 0.248 |
|  | CI | -474.33 – -277.29 | 0.36 – 1.05 | -0.30 – 0.40 | 0.02 – 0.03 |  |  |  |  |  |
|  | p-value | **<0.001** | **0.003** | 0.715 | **<0.001** |  |  |  |  |  |
|  | df | 258.80 | 6.11 | 5.60 | 258.81 |  |  |  |  |  |
| DHA/EPA | Estimates | 341.14 | -0.62 | 0.29 | -0.02 | 1.00 | 0.00 | 9 | 264 | 0.222 |
|  | CI | 236.57 – 445.71 | -0.99 – -0.25 | -0.08 – 0.66 | -0.02 – -0.01 |  |  |  |  |  |
|  | p-value | **<0.001** | **0.006** | 0.104 | **<0.001** |  |  |  |  |  |
|  | df | 258.80 | 6.11 | 5.60 | 258.81 |  |  |  |  |  |
| ALA | Estimates | -102.60 | -2.54 | -2.29 | 0.01 | 1.41 | 0.00 | 9 | 264 | 0.484 |
|  | CI | -227.03 – 21.84 | -2.98 – -2.09 | -2.74 – -1.85 | -0.00 – 0.01 |  |  |  |  |  |
|  | p-value | 0.106 | **<0.001** | **<0.001** | 0.094 |  |  |  |  |  |
|  | df | 258.80 | 6.11 | 5.60 | 258.81 |  |  |  |  |  |

**Table S3.** PERMANOVA of fatty acids of PHASE (experimental feeding Phase) of fish permutations = 999, method = Bray Curtis

|  | Df | Sum of Sqs | R2 | F | p-value |
| --- | --- | --- | --- | --- | --- |
| **Treatment** | 2 | 0.0765 | 0.4333 | 45.112 | 0.001 |
| **Residual** | 118 | 0.1001 | 0.5667 |  |  |
| **Total** | 120 | 0.1766 | 1.0000 |  |  |

**Table S4.**
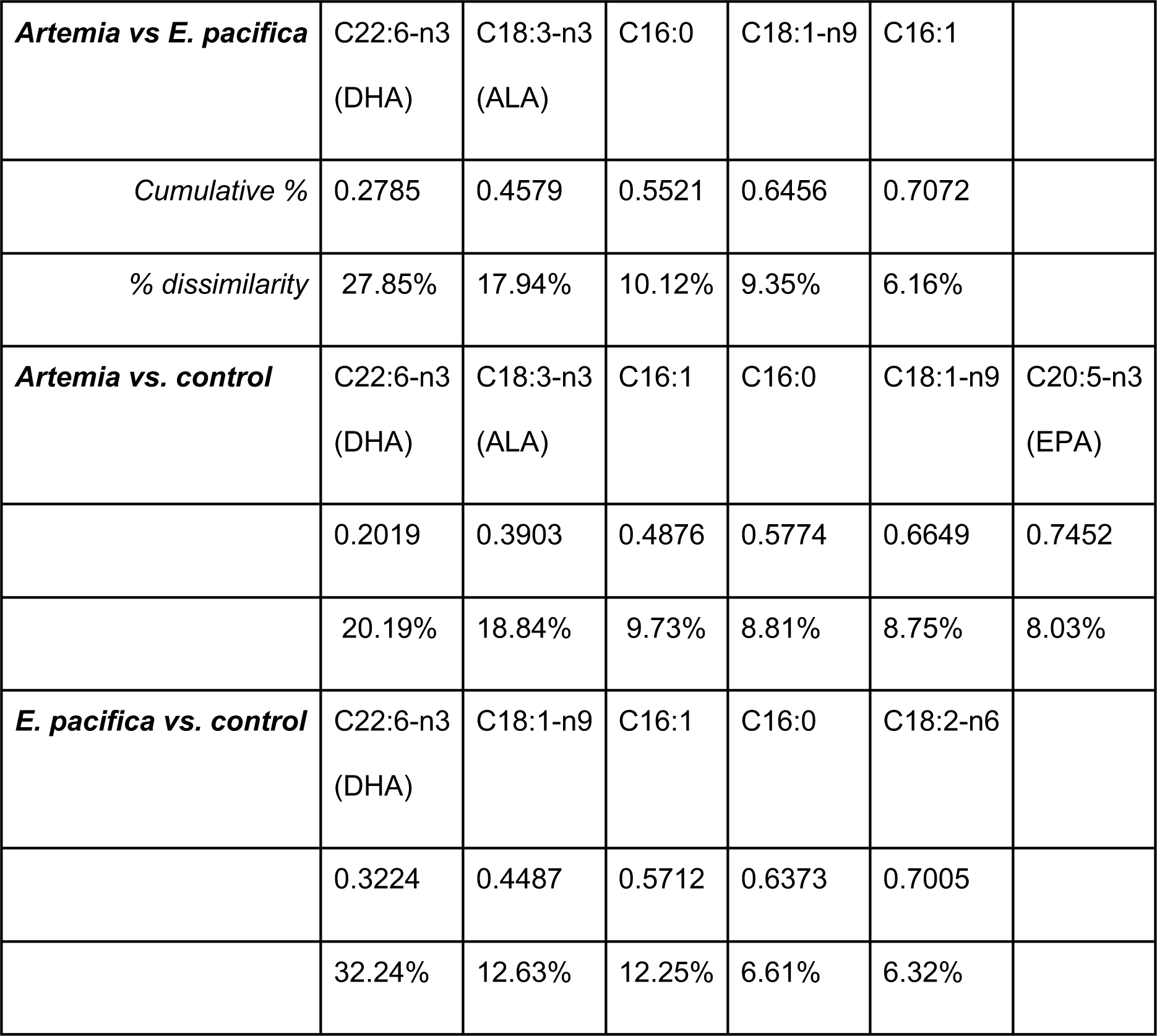
SIMPER Cumulative contributions of most influential fatty acids.

**Figure S1.**
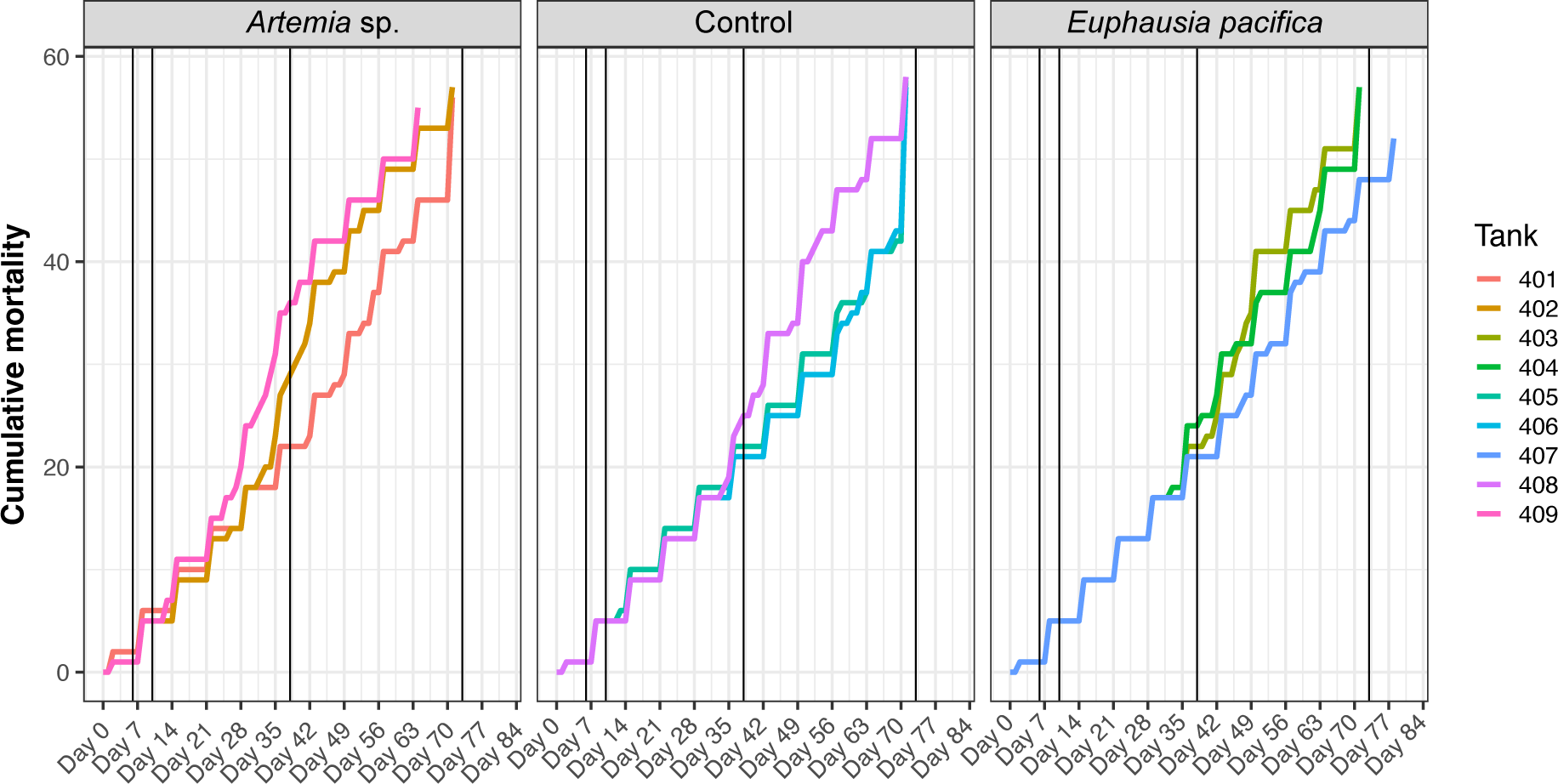
Cumulative mortality of fish in the different treatments (*Artemia* sp., control, *Euphausia pacifica*) over time.

## Notes

### Competing Interest Statement

The authors have declared no competing interest.

### Summary of Updates

The changed the focus of the manuscript to a more global perspective how future predicted reduction in phytoplankton DHA and EPA will occur and how this would affect salmon

